# Fission Yeast Cells Grow Approximately Exponentially

**DOI:** 10.1101/489708

**Authors:** Mary Pickering, Lauren Nicole Hollis, Edridge D’Souza, Nicholas Rhind

**Affiliations:** Biochemistry and Molecular Pharmacology, University of Massachusetts Medical School, Worcester MA 01605 USA

**Keywords:** Fission Yeast, Schizosaccharomyces pombe, Cell Growth Kinetics, Cell Size, Cell Cycle

## Abstract

How the rate of cell growth is influenced by cell size is a fundamental question of cell biology. The simple model that cell growth is proportional to cell size, based on the proposition that larger cells have proportionally greater synthetic capacity than smaller cells, leads to the predication that the rate of cell growth increases exponentially with cell size. However, other modes of cell growth, including bilinear growth, have been reported. The distinction between exponential and bilinear growth has been explored in particular detail in the fission yeast *Schizosaccharomyces pombe*. We have revisited the mode of fission yeast cell growth using high-resolution time-lapse microscopy and find, as previously reported, that these two growth models are difficult to distinguish both because of the similarity in shapes between exponential and bilinear curves over the two-fold change in length of a normal cell cycle and because of the substantial biological and experimental noise inherent to these experiments. Therefore, we contrived to have cells grow more than two fold, by holding them in G2 for up to eight hours. Over this extended growth period, in which cells grow up to 5.5-fold, the two growth models diverge to the point that we can confidently exclude bilinear growth as a general model for fission yeast growth. Although the growth we observe is clearly more complicated than predicted by simple exponential growth, we find that exponential growth is a robust approximation of fission yeast growth, both during an unperturbed cell cycle and during extended periods of growth.

## INTRODUCTION

The relationship between the rate of cellular growth and cell size has critical implications for the maintenance of cell-size homeostasis. If cells grow at a rate independent of cell size, size homeostasis can be maintained by growing for a fixed amount of time each cell cycle (Brooks, 1981; Fantes and Nurse, 1981; Conlon and Raff, 2003). However, if growth is proportional to size — that is, if big cells grow faster than small cells — big cells will tend to get bigger and small cells will tend to stay smaller, necessitating an active size control mechanism to maintain size homeostasis in a population (Amodeo and Skotheim, 2015). Therefore, in order to understand how cells regulate their size, it is essential to understand how their growth rate changes with size.

The natural postulate that growth rate is proportional to cell size follows from the assumption that, absent developmental or other constraints, cells grow as quickly as possible. Since a cell’s biosynthetic capacity in nutrient-rich growth conditions is largely limited by its translational capacity, larger cells, with more ribosomes, should grow faster than smaller cells. A simple corollary of this postulate is that cell mass increases exponentially over time. Exponential cellular growth has been reported for many cell types, both prokaryotic and eukaryotic (Schaechter et al., 1958; Fantes and Nurse, 1977; Johnston et al., 1977; Cooper, 1988; Dolznig et al., 2004; Sung et al., 2013; Cermak et al., 2016). However, there have been a number of reports of non-exponential cell growth, as well (Prescott, 1955; Mitchison and Nurse, 1985; Conlon and Raff, 2003; Son et al., 2012). Thus, it is unclear if exponential growth is the general situation from which cells sometimes deviate, or if it is one of many possible modes of cellular growth.

Growth kinetics of the fission yeast *Schizosaccharomyces pombe* have been particularly controversial (Cooper, 1998; Mitchison et al., 1998; Sveiczer et al., 2014). Fission yeast cells were initially observed to grow approximately exponentially in length (Mitchison, 1957). However subsequent work from the same lab reported bilinear growth—that is, growth at a constant, size-independent rate until a discrete rate change point (RCP) midway through the cell cycle at which point cell growth rate increases (Mitchison and Nurse, 1985). Bilinear growth has also been reported by three more recent papers (Sveiczer et al., 1996; Baumgartner and Tolic-Norrelykke, 2009; Horvath et al., 2013). However, these conclusions have been challenged by the claim that the data presented is equally consistent with exponential growth (Cooper, 1998; Cooper, 2013). A fundamental issue in this controversy is that the expected difference between exponential and bilinear growth is subtle; the maximum expected difference between an exponential and a bilinear growth curve over a normal cell doubling is only about 3% (Figure S1), less than the experimental error in most growth-kinetics experiments (Buchwald and Sveiczer, 2006; Cooper, 2013). Several studies have determined which models are better statistical fits to various datasets and found that a bilinear model generally fits better than an exponential one (Buchwald and Sveiczer, 2006; Baumgartner and Tolic-Norrelykke, 2009). Nonetheless, the difference in goodness-of-fit between the two models is not sufficient to exclude either one (Buchwald and Sveiczer, 2006).

We have revisited the question of fission yeast growth kinetics using high-resolution video microscopy. Our results with unperturbed cells appear consistent with exponential growth. However, because, as previously noted (Buchwald and Sveiczer, 2006), the predictions of exponential and bilinear growth are quite similar over the two-fold growth of unperturbed cells, we cannot, using that data alone, exclude the bilinear hypothesis. Therefore, we also examined the growth kinetics of cells held in G2, which grow to be much longer than twice their birth length. These cells clearly show a size-dependent increase in growth rate incompatible with bilinear growth. Although actual cellular growth kinetics are clearly more complicated than a simple exponential model, our results suggest that an exponential model is a robust approximation for fission yeast growth kinetics over normal and extended cell growth.

## RESULTS

### Average Cell Growth is Proportional to Cell Size Across a Wide Range of Cell Sizes

In a normal cell cycle, cell mass must double prior to division into two daughter cells. A simple prediction of exponential growth is that the amount of time required for this doubling is independent of cell size. Large cells require more growth than small cells in order to double in size; however, if growth rate is proportional to size (and thus exponential), then the faster growth of larger cells balances the larger amount of growth required to double, resulting in equal doubling times for large and small cells. To test if this prediction is met in fission yeast, we grew populations of isogenic cells under conditons that resulted in different distributions of cell sizes. Specifically, we varied the size of cells in asynchronous culture by regulating the expression of the Wee1 mitotic inhibitor from a promoter regulated by ZEV, a synthetic, estradiol-responsive transcription factor (Ohira et al., 2017). By varying the dose of β-estradiol from 0 to 100 nM, we obtained cultures that vary in length at division from 13.6 to 31.3 *μ*m (Figure 1). Despite this 2.3-fold increase in cell size, the doubling times for these cultures varied by only about 15% (147±26 minutes) and the variation did not correlate with size. Wild-type control cultures, and control cultures of a strain that expresses the ZEV transcription factor but does not overexpress Wee1, varied in the same range (137±19 minutes). Finally, we also measured the doubling time of *wee1-50ts* cells at the semi-restrictive temperature of 30°C. These cells divide at 9.1±0.5 *μ*m, extending our size range to 3.4 fold, but still double at a similar rate (137±27 minutes). These results are consistent with previous observations that the doubling time of cells is independent of cell size (Russell and Nurse, 1987; Sveiczer et al., 1996; Zhurinsky et al., 2010), indicating that smaller cells gain mass more slowly and larger cells gain mass faster, resulting in cells that have consistent growth kinetics overall.

**Figure 1:**
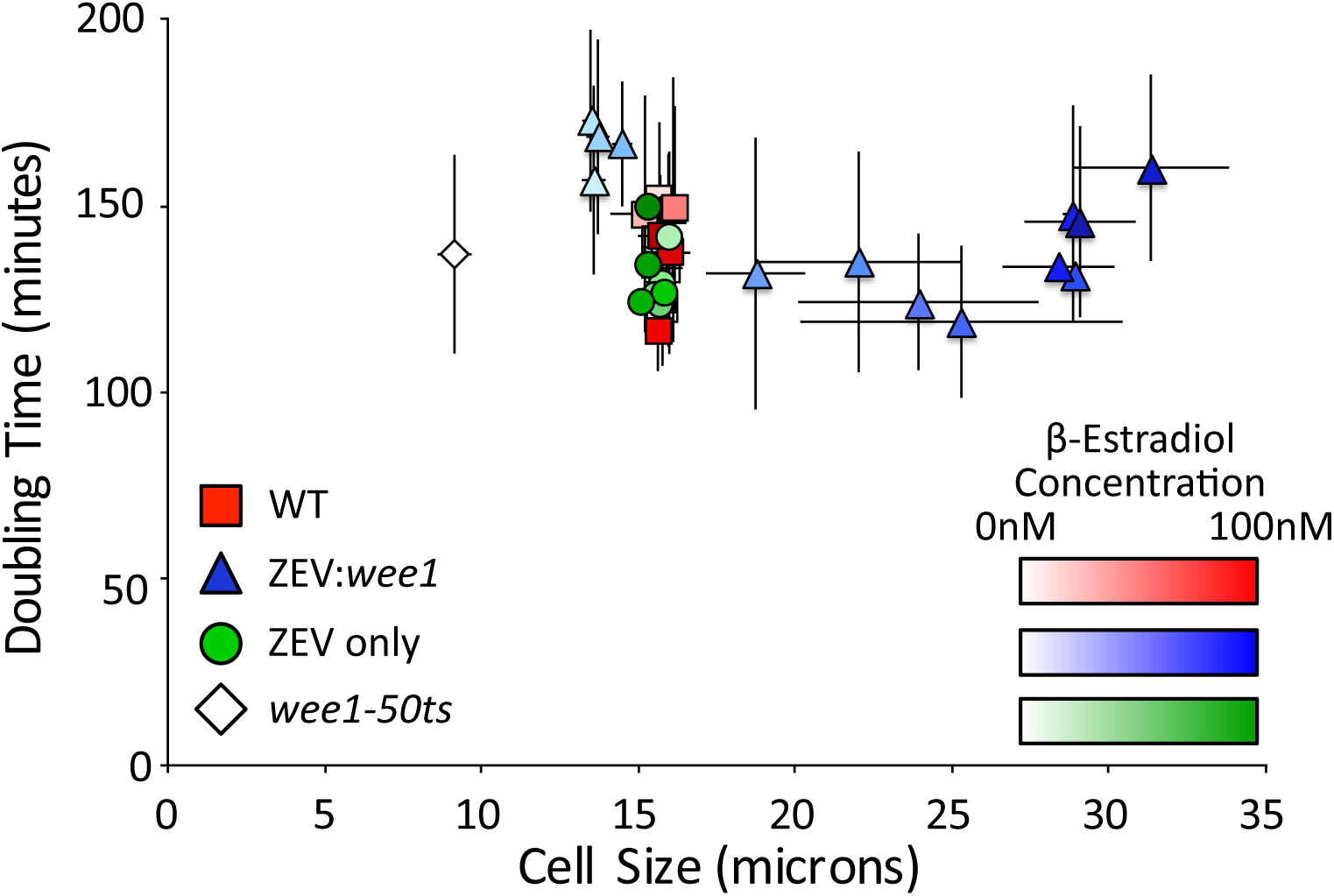
The doubling time of cell populations is independent of cell size. The optical densities (OD) at 600 nm of asynchronous cultures of wild-type (yFS105), *wee1-50* (yFS131), *ZEV:wee1* (yFS970) and *ZEV* control (yFS949) cells were measured over time. Where indicated, cells were treated with β-estradiol at 0.1, 0.31, 1.0, 3.1, 5.0, 6.0, 6.5, 7.0, 7.5, 10, 31, or 100 nM. The doubling time of the culture was calculated using the exponential rate from the sigmoidal fit of the data. The lengths of at least 50 septated cells per treatment were measured to calculate average length at septation.

The fact that cell doubling times are independent of cell size demonstrates that the rate of cell growth, averaged over the cell cycle, is proportional to cell size. This observation rules out the possibility that cells grow with a fixed linear rate. However, it does not exclude the possibility that they grow at a linear rate proportional to birth size, or that they grow bilinearly, with either the growth rates or the position of the RCP being size dependent. Therefore, to determine the growth kinetics of individual cells, we assayed growth kinetics by high-resolution video microscopy.

### Wild-type Cells Grow Approximately Exponentially

To determine the mode of individual cell growth, we used time lapse microscopy to record the length of cells growing in a microfluidic chamber. Initially, we examined 12 wild-type cells (Figure 2). Birth length of the cells varied from about 7 to 10 *μ*m and cells approximately doubled in length as they progressed to septation (Figure 2A,E). Cells grow for about 3/4 of the cell cycle and then enter a constant-length phase during which nuclear and cellular division occur. By plotting normalized cell lengths, it is apparent that there is substantial variation in the fold growth over the cell cycle (from 1.6 to 2.2 fold) and the growth rate (from doubling times of 169 to 263 minutes, Figure 2E). Such heterogeneity in individual-cell growth parameters requires an active size-control mechanism to maintain size homeostasis at the population level (Amodeo and Skotheim, 2015).

**Figure 2:**
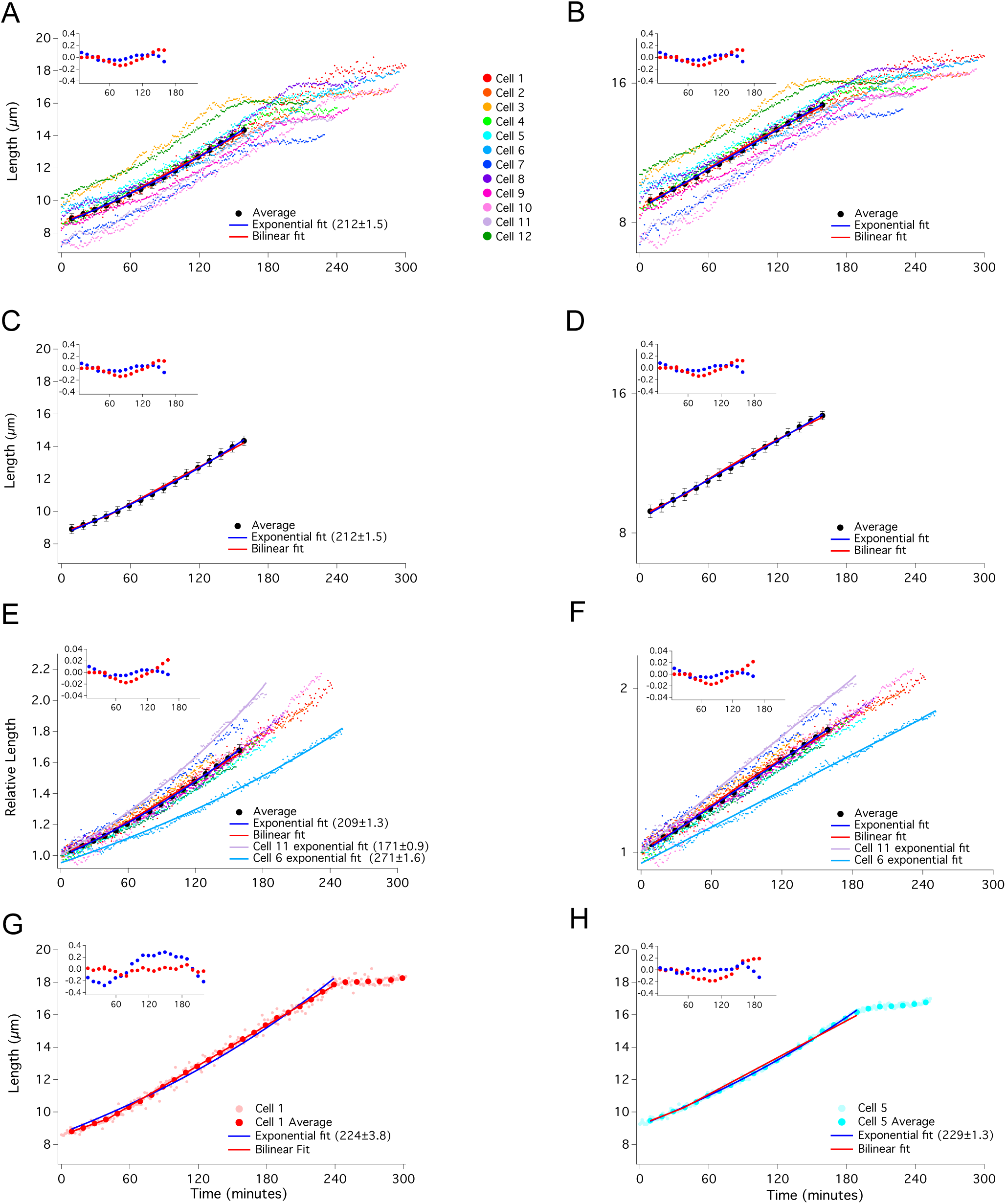
Growth of wild-type cells is well fit by both bilinear and exponential curves. Lengths of twelve individual wild-type (yFS105) cells were recorded at 1 minute intervals from birth to septation and plotted on a linear (A) or log (B) scale. The average of the cells in 10 minute windows ± standard error of the mean, is shown on a linear (C) and log (D) scale. Exponential fit (with doubling time in parentheses), bilinear fit, and residual plot for these fits (inset) are shown. Birth lengths of cells were normalized to 1 and plotted using a linear scale (E) or a log-scale (F). The individual and average data points for a cell exhibiting an apparently bilinear (G) or an apparently exponential (H) mode of growth are shown.

By visual inspection, some of the growth curves look exponential, some bilinear, and others have more complicated shapes (Figure S2), as previously observed (Horváth et al., 2016). To demonstrate the heterogeneity of growth curve shapes, we fit exponential and bilinear curves to two cells representative of each shape (Figure 2G,H). We also calculated the average growth trajectory of all 12 cells and fit it to both an exponential and bilinear curve (Figures 2A-D). Both curves fit the raw and normalized measurements of cell lengths within the experimental variation (Figures 2A-F). Thus, for reasons presented in the Discussion, we conclude that fission yeast cell growth can accurately be described as approximately exponential.

Throughout this work we refrain from using statistical methods to quantitatively compare the goodness of fit of specific exponential or bilinear curves to our data. We agree with the published conclusion that, over a two-fold change in length, such methods cannot distinguish between the two growth models (Buchwald and Sveiczer, 2006) and will claim below that over larger changes in length, the difference is obvious. To make the difference between the data and the model curves easier to compare, we include an inset of the residuals of the fits for each data set to which we fit a model curve. To quantitate the difference between the data and the fit, we calculated the root-mean-squared difference (RMSD, a simple statistic for the difference between two datasets that captures the average difference across the range of data points). For the bilinear fit, the RMSD is 0.08 *μ*m; for the exponential fit is is 0.05 *μ*m. Both of these well within the average standard error of the average growth trajectory, which is 0.31 *μ*m.

### Cells Grow Approximately Exponentially when G2 Growth is Extended

Exponential and bilinear modes of growth are similar over small fold changes in length, such as the doubling of cell length during the cell cycle. However, over larger fold changes in length, the two growth models diverge dramatically. Whereas the maximum difference between the two growth models over a two-fold change in length is less than 3%, extending the growth to four-fold reveals a greater than 20% difference (Figure S1). To take advantage of this divergence, we held cells in G2, allowing cells to grow much longer than two-fold in length. Initially, we took advantage of the *ZEV:wee1* overexpression system. Asynchronous, untreated *ZEV:wee1* cells, which divide at about 9 *μ*m, were treated with 31 nM β-estradiol and immediately subjected to time lapse microscopy to capture cells as their lengths increased from birth to septation, which in 31 nM β-estradiol is about 27 *μ*m. We were thus able to observe growth over an extended range, with some cells growing almost four-fold in length (Figure 3A). The individual growth curves look approximately exponential, although there is plenty of deviation from simple exponential growth. Cell 14, for instance, has a number of brief pauses in growth (Figure S3). The approximately exponential nature of the growth curves is particularly clear when the data is plotted on a semi-log plot (Figure 3C). Some of the curves are straight, indicative of exponential growth, but some curves, and the average of all of the curves, are slightly sub-exponential, suggesting that larger cells may grow somewhat more slowly than predicted by a simple exponential model (Figures 3A-F and S3). When a bilinear curve is fit to individual-cell or averaged data, it fits reasonably well for the first doubling of cell size, but diverges as the cells continue to grow proportionally with size (Figure 3). For the bilinear fits, we placed the RCP close to 60 minutes, similar to its position in our wild-type data and to the RCP reported in other studies (Baumgartner and Tolic-Norrelykke, 2009; Horvath et al., 2013). The deviation of the bilinear fit from the data is apparent in the residuals of the fits to individual cells and to the average growth trajectory (Figures 3,S3). The RMSD for the bilinear fit (0.48 *μ*m) is more the twice that of the exponential fit (0.21 *μ*m), although both are less that the average of the standard error of the average growth trajectory (0.81 *μ*m).

**Figure 3:**
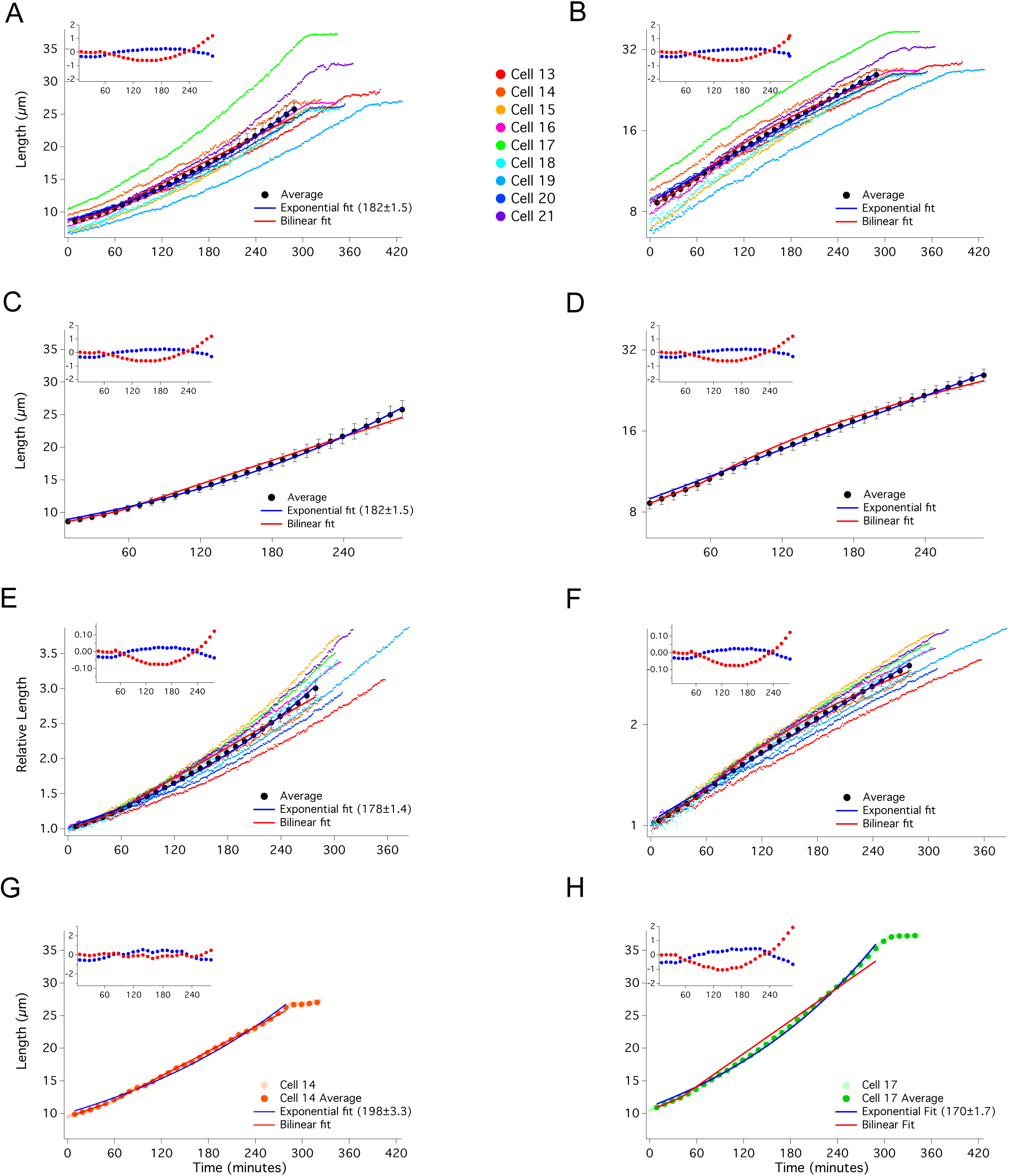
Extended cell growth is well fit by an exponential curve. Asynchronous, *ZEV:wee1* expressing (yFS970) cells were treated with 31 nM β-estradiol to elongate growth in G2. Cell lengths were recorded at 1 minute intervals from birth to septation and plotted on a linear (A) or log (B) scale, as in Figure 2. The average of the cells in 10 minute windows ± standard error of the mean, is shown on a linear (C) and log (D) scale. Birth lengths of cells were normalized to 1 and plotted using a linear scale (E) or a log-scale (F). The individual and average data points for a cell exhibiting an apparently bilinear (G) or an apparently exponential (H) mode of growth are shown.

To arrest cells in G2 for even longer and thus get an even larger divergence between the exponential and bilinear models, we used the stronger *adh1:wee1-50* overexpression system. These cells constitutively overexpress the temperature-sensitive Wee1-50 protein. At the restrictive temperature of 35°C, cells are born at about 5 *μ*m but arrest in G2 when shifted to 25°C (Russell and Nurse, 1987). We shifted asynchronous cells to 25°C and followed them by time lapse microscopy for 16 hours. For incubations longer than 8 hours, the cells stop growing (Figure S4). However, since all the cells stop growing at the same time (between 8 and 10 hours) and not at the same length (terminal lengths vary between about 30 and 55 *μ*m), we believe the cessation of growth is due to the depletion of some growth-limiting nutrient or the accumulation of some toxic waste product within the microfluidic chamber, not any intrinsic limitation to fission yeast growth. Similar to the *ZEV:wee1* cells, these cells grew approximately exponentially, with a number of cells deviating from a simple exponential curve (Figures 4A-F, S5). An exponential model fits both the individual cell and averaged cell data over its full extent. The RMSD for the exponential fit is 0.31 *μ*m, well within the standard error of 0.87 *μ*m. In contrast, a bilinear model fits the data only for the initial doubling of cell size, after that it quickly diverges, consistent with cells growing proportionally with size across the entire time course. The RMSD of the bilinear fit is 1.1 *μ*m, larger that the average standard error of 0.87 *μ*m. Furthermore, the bilinear fit deviates beyond the standard error of the average growth trajectory at specific timepoint, from 140 minutes to 300 minutes and after 440 minutes (Figure 4C).

**Figure 4:**
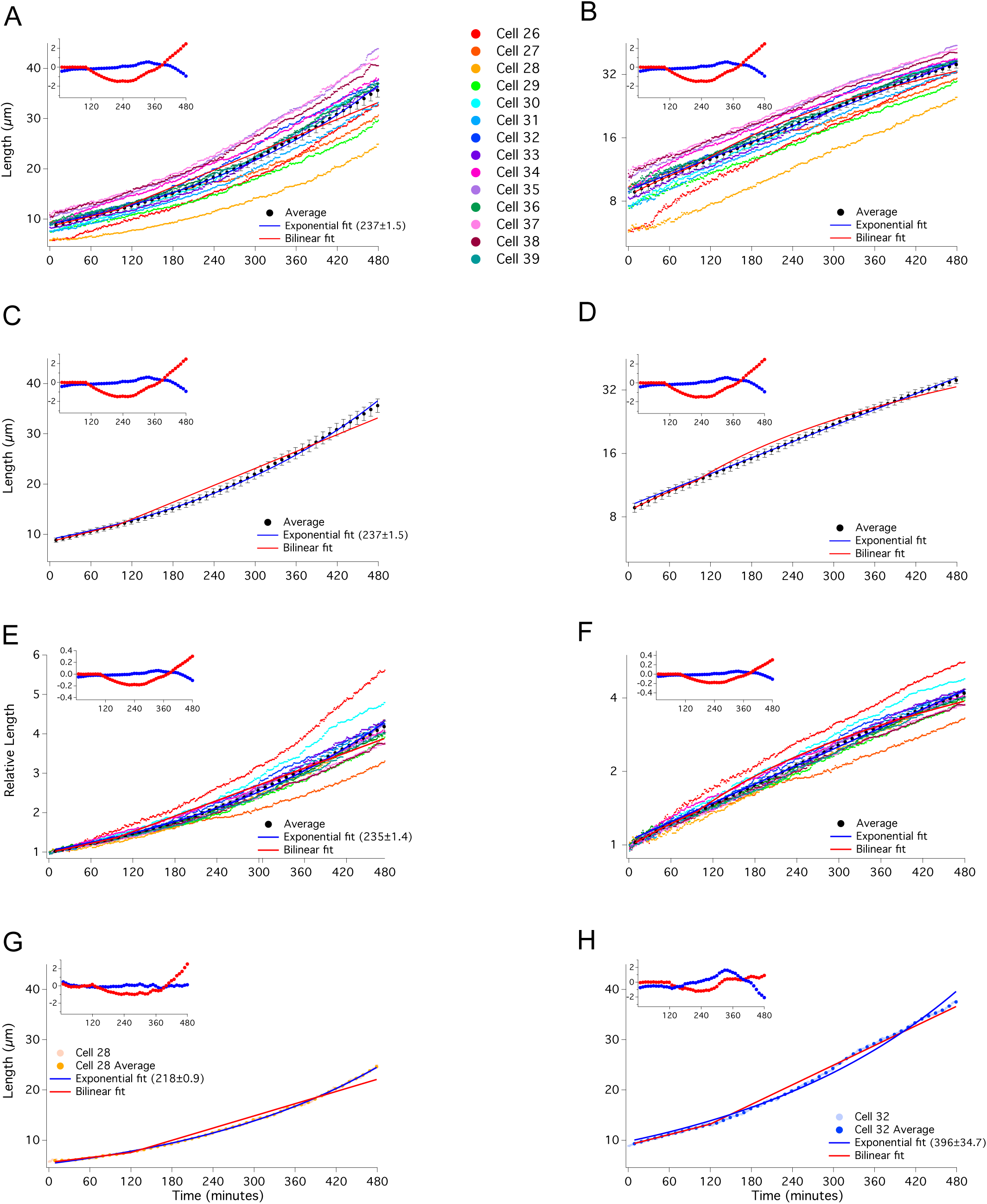
Extended cell growth is well fit by an exponential curve. Asynchronous, *adh1:wee1-50* (yFS145) cells grown at 35°C were shifted to 25°C to elongate growth in G2. Cell lengths were recorded at 1 minute intervals and data were plotted on a linear (A) or log (B) scale, as in Figure 2. The average of the cells in 10 minute windows ± standard error of the mean, is shown on a linear (C) and log (D) scale. Birth lengths of cells were normalized to 1 and plotted using a linear scale (E) or a log-scale (F). The individual and average data points for a cell exhibiting an apparently exponential (G) or an apparently bilinear (H) mode of growth are shown.

## DISCUSSION

A simplifying assumption in many models of cell size control is that cell growth is exponential (Fantes et al., 1975; Amodeo and Skotheim, 2015), which in turn is based on the assumption that, for cells growing in a rich medium, growth rate is limited by protein synthetic capacity and therefore by the number of ribosomes. It follows from this assumption that growth rate should be proportional to cell size, which correlates closely with ribosome number (Bakshi et al., 2012), and is thus exponential. Fission yeast is an excellent system in which to study cell growth rate because its cylindrical cells grow only by elongation, making growth easy to measure microscopically (Mitchison, 1957). Many groups have taken advantage of this property to measure the growth kinetics of single fission yeast cells by time-lapse microscopy (Mitchison, 1957; Mitchison and Nurse, 1985; Miyata et al., 1988; Sveiczer et al., 1996; Baumgartner and Tolic-Norrelykke, 2009; Horvath et al., 2013; Horváth et al., 2016). Although fission yeast growth kinetics have been reported by several groups to be bilinear, reanalysis of some of the same data has led to the conclusion that growth is exponential (Mitchison and Nurse, 1985; Sveiczer et al., 1996; Cooper, 1998; Baumgartner and Tolic-Norrelykke, 2009; Cooper, 2013). However, both biological variation in growth patterns and experimental noise result in heterogeneous data, in which different cells display different patterns of growth (Figures S2, S3, S5 and Horváth et al., 2016). Sophisticated curve-fitting approaches have been used to determine which model best fits the data, but, although bilinear curves fit marginally better than exponential curves, neither fit can be excluded with the available data. (Buchwald and Sveiczer, 2006; Baumgartner and Tolic-Norrelykke, 2009). The problem is that, over the two-fold change in length of a wild-type cell doubling, the maximum difference between an exponential and bilinear curve is only about 3%, which is within experimental variation (Figure S1). Our data for the growth of unperturbed wild-type cells is consistent with previous work, showing reasonable fits with both exponential and bilinear curves (Figure 2).

Although exponential and bilinear curves are similar within a two-fold length range, they diverge quickly with longer growth, reaching over 20% difference after a four-fold change in length (Figure S1). Therefore, we held cells in extended G2 by over expressing the Wee1 mitotic inhibitor in two different ways (Figures 3,4). These treatments allowed cells to grow up to 5.5 fold in length. Over these time courses, cell growth is still reasonably fit by an exponential curve, but clearly deviates from a bilinear curve (Figures 3,4). These results parallel those done by arresting cells with a temperature sensitive *cdc2* allele (Miyata et al., 1988). In that case, the growth curves were described as multi-linear with up to three RCPs; however, it was noted that such multi-linear curves produce pseudo-exponential growth kinetics (Miyata et al., 1988). Although these patterns of extended growth do not prove that fission yeast growth is exponential, they do falsify the hypothesis that fission yeast growth can generally be described as bilinear.

The question of whether the extended growth of fission yeast held in G2 is better described as exponential or multi-linear prompts consideration of the purpose of fitting a mathematical model to experimental data. If the model is predicted by a biological hypothesis — such as the prediction of exponential growth from the hypothesis that larger cells grow proportionally faster than smaller cells because they have proportionally more ribosomes — curve fitting can be used to test that hypothesis. A failure of an exponential curve to fit the data would falsify the hypothesis and demonstrate that some other mechanism of cell growth regulation must exist. Since our data (Figures 2-4), and previously published data (Buchwald and Sveiczer, 2006; Cooper, 2013), is reasonably well fit by an exponential curve, it is compatible with the proportional-growth hypothesis.

If a mathematical model is fit to data empirically, without a biological rational, the model can be used to generate a hypothesis. The bilinear fit to the fission yeast growth data generates the hypothesis that there is a physiological change at the RCP that allows the cells to grow faster (Sveiczer et al., 1996). Several cell cycle events, such as DNA replication and the switch from growth at one cell tip to growth at both cell tips, known as new-end take off (NETO), have been suggested as the cause of the fission yeast RCP, but none have been experimentally confirmed (Sveiczer et al., 1996; Baumgartner and Tolic-Norrelykke, 2009). Although it is plausible that a cell cycle event could affect the cellular growth rate, it is more difficult to image how multiple RCPs could be explained in a single G2, making it harder to generate hypotheses compatible with the multi-linear model. For that reason, we prefer the exponential model over the multi-linear model.

Another way to think about the question of growth-rate kinetics is to wonder how cellular growth could not be exponential. In Figure 1, we show that, averaged over the cell cycle, larger cells grow proportionally faster than smaller cells. If cells grow exponentially throughout the cell cycle, such average proportionality is ensured. However, if cells grow in one or more linear segments, there must be some mechanism that prevents larger cells from growing proportionally faster, until an RCP is reached. One plausible hypothesis for the cause of an RCP is the doubling of the DNA during replication, which would double the number of transcriptional templates, plausibly increasing the rate of transcription (Baumgartner and Tolic-Norrelykke, 2009). However, the RCP reported during wild-type fission yeast growth does not correlate with the timing of DNA replication (Sveiczer et al., 1996; Baumgartner and Tolic-Norrelykke, 2009). Furthermore, transcription has been shown to scale with cell size, not DNA copy number, within the normal range of cell size (Zhurinsky et al., 2010; Marguerat and Bahler, 2012). Finally, DNA replication cannot explain multiple RCPs during extended G2 growth. Another cell cycle event proposed to explain the RCP is NETO (Mitchison and Nurse, 1985). However, NETO has been shown to not correlate with the RCP (Sveiczer et al., 1996; Baumgartner and Tolic-Norrelykke, 2009), nor could it explain multiple RCPs. Finally, even if a cell cycle event could lead to an RCP, it is unclear how the position of the RCP in the cell cycle and the growth rates of the linear segments would be coordinated to ensure average proportional growth. Given the lack of a plausible explanation for bi- or multi-linear growth, and the natural prediction of exponential growth, we favor the conclusion that fission yeast growth kinetics are adequately described by an exponential model.

Although we conclude that exponential growth is an adequate description of fission yeast growth kinetics, the growth patterns of individual cells clearly deviate from a simple exponential curve (Figures S2,S3,S5). Much of the deviation from exponential growth appears to be unpredictable heterogeneity, such as the early pause in growth observed in Cell 11 and later discontinuities in growth in Cells 08 and 14. This heterogeneity we ascribe to varying environmental circumstances or physiological perturbations that cause individual cells to deviate from what would otherwise be exponential growth. In particular, it is possible NETO can, in some cell growth trajectories, cause a discontinuity of growth in length. In addition to what appears to be unpredictable heterogeneity, we note that many cells held in G2 grow slightly slower than predicted by a simple exponential model as they elongate (Figures 3,4). This sub-exponential growth may indicate that larger cells grow less efficiently than smaller cells, perhaps due to reduced relative transcriptional capacity as cells increase in size (Zhurinsky et al., 2010). However, it is also possible that this effect is due to growth in the microfluidic chambers we used and that cells in optimal growth conditions would more closely approximate exponential growth. For these reasons, we do not claim that fission yeast growth is strictly exponential. Instead, we propose that fission yeast growth is approximately exponential and that no specific growth control mechanisms are required to explain non-exponential growth.

## MATERIALS AND METHODS

### Yeast strains and growth conditions

The *Schizosaccharomyces pombe* strains used in this study were yFS105 (*h− leu1-32 ura4-D18*, lab stock), yFS131 (*h*+ *leu1-32 ura4-? wee1-50*, lab stock), yFS145 (h+ *leu1-32 ura4-D18 wee1::pWAU-50(adh1:wee1-50 ura4)* (Russell and Nurse, 1987)), yFS949 *(h− leu1-32::pFS461(adh1:ZEV leu1) ura4-D18 ade6-210, his7-366*, (Ohira et al., 2017)) and yFS970 *(h− leu1-32::pFS461(adh1:ZEV leu1) ura4-D18 ZEVpr:wee1::(kanMX6)*, (Ohira et al., 2017)). Fission yeast cultures were grown in YES media at either 25°C (yFS131) or 30°C (all other strains), as previously described (Forsburg and Rhind, 2006). When indicated, β-estradiol (E2758, Sigma Aldrich) dissolved in ethanol at 10 mM was added to the indicated final concentration.

### Doubling time analysis and cell length measurements

Cells were grown to mid log phase, diluted to optical density (OD) 0.02, and plated in triplicate in a 96-well microtiter plate. For samples treated with β-estradiol, cells were treated for at least 8 hours in the indicated concentration prior to dilution. The samples were grown at 30°C with shaking for 36 hours in a Biotek Eon Microplate Spectrophotometer and the OD 600nm of each well was recorded at 5 min intervals. To determine the doubling time of the cultures, sigmoidal curve fitting analysis was calculated in Igor Pro 6.0 (Wavemetrics). Cells from these cultures were photographed using DIC optics on a Zeiss Axioshop 2.0. The cell lengths for at least 50 cells per treatment were measured in Image J (Schneider et al., 2012).

### Individual cell growth analysis

Using the Freestyle Fluidics approach (Walsh et al., 2017), time-lapse movies of individual cells were generated on a DeltaVision OMX with DIC optics. Cells were grown to mid log and were then diluted to OD 0.04, left untreated or treated with 31 nM β-estradiol, and immediately spotted onto a lectin-coated (L1395, Sigma Aldrich) glass-bottomed 35mm dish (P35G-1.5-14-C MatTek) under 1 cm of FC40 oil (gift of Peter Cook). Images were recorded in one-minute intervals for 16 hours. The *adh1:wee1-50* cells were grown for at least 16 hours at 35°C prior to diluting to OD 0.04 and were shifted to 25°C in a temperature-controlled chamber for the duration of the time-lapse image collection. The length of individual cells was measured in every frame from birth to septation using Image J (Schneider et al., 2012). All cell lengths are available in Table S1. To smooth the curves, the cell length was averaged every 10 minutes in overlapping 20 minute windows. For each dataset, the value in each window for all cells was averaged and standard error was calculated. Data was plotted in Igor Pro and fit using the CurveFit function to either an exponential (y=(2a)^^^(tx), where a is the initial length and t is the doubling time) or bilinear (y=a_1_+b_1_x for x≤RCP and y=a_2_+b_2_x for x≥RCP) equation.

## Supporting information

## ACKNOWLEDGEMENTS

We gratefully acknowledge Christiana Baer and the Sanderson Center for Optical Experimentation (SCOpE) for access to, and technical support for, the DeltaVision OMX microscope, Peter Cook for sharing his Freestyle Fluidics technology before publication, and Peter Pryciak for helpful comments on a draft the manuscript.

---

**Figure S1:**
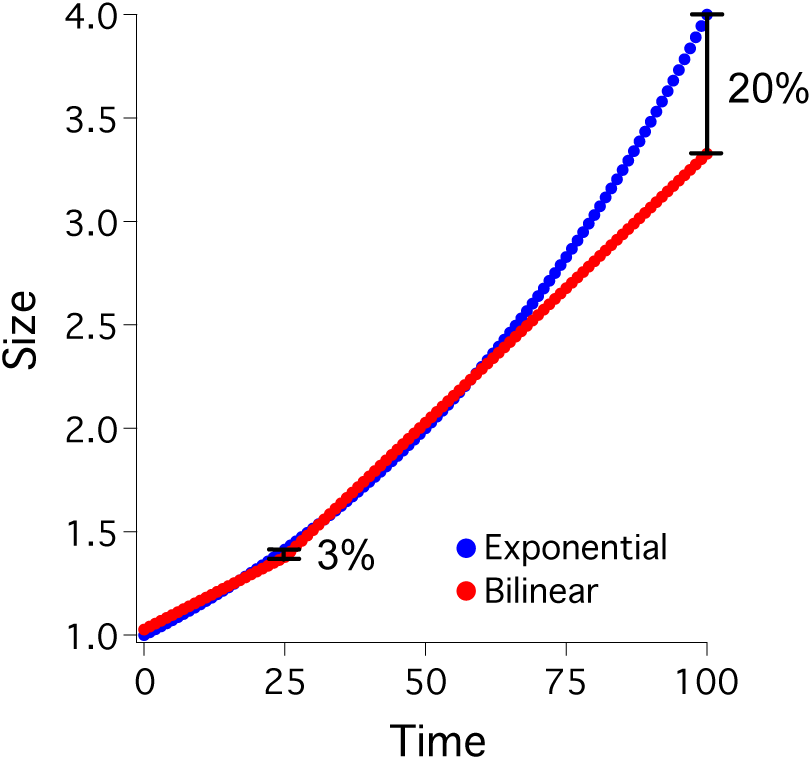
Ideal exponential and bilinear growth curves. Graphical representation of ideal growth curves exhibiting exponential or bilinear growth over two doublings in length is shown. The curves are superimposed so as to minimize their difference over the first doubling. The difference between the two curves at 25 minutes is 2.7%; the difference at 100 minutes is 20.2%.

**Figure S2:**
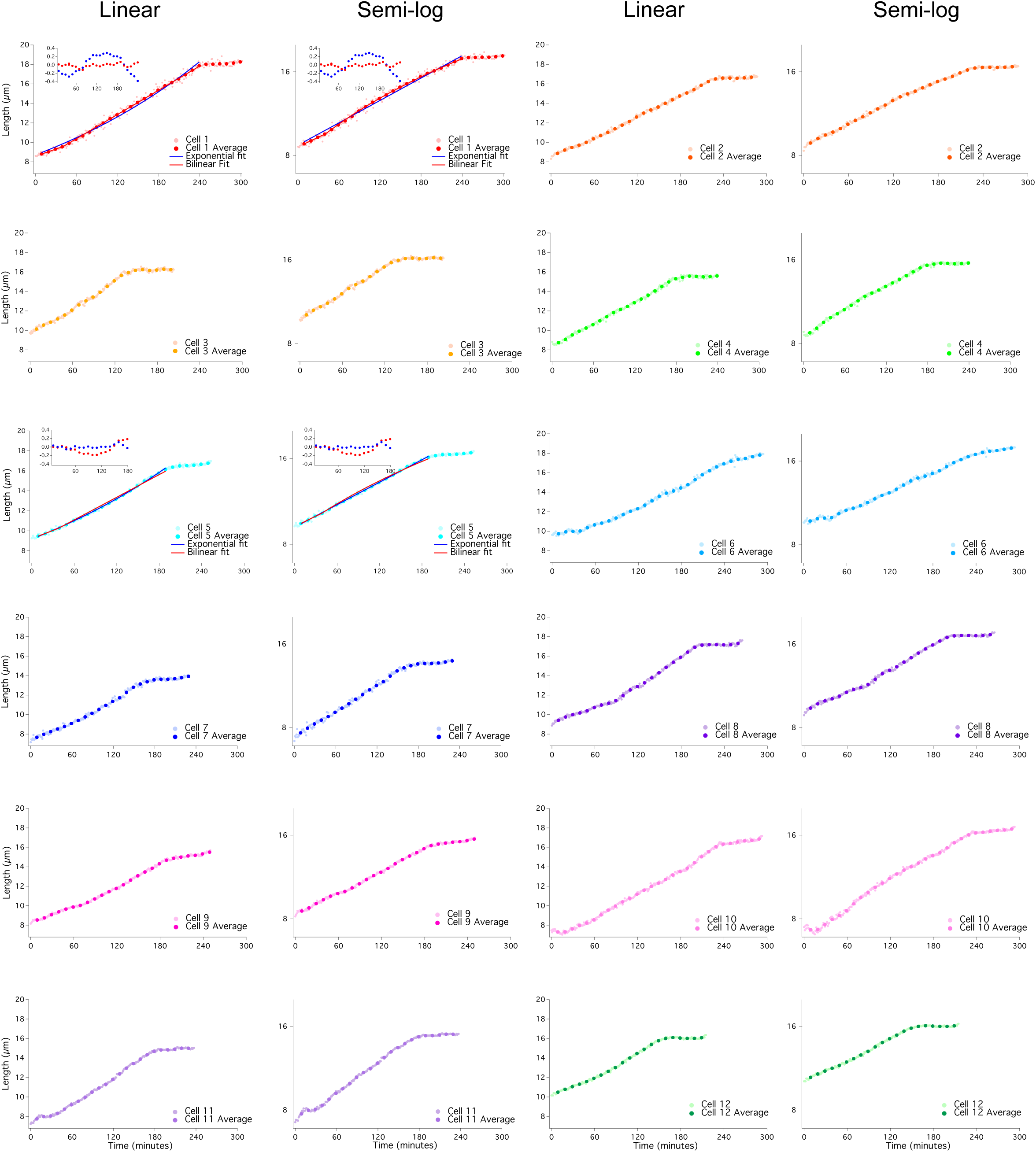
Individual growth data for 12 wild-type cells. Lengths of twelve individual wild-type (yFS105) cells were recorded at 1 minute intervals from birth to septation and plotted on a linear or log scale. Cell length was averaged every 10 minutes in overlapping 20 minute windows. Fits from Figure 2 are included.

**Figure S3:**
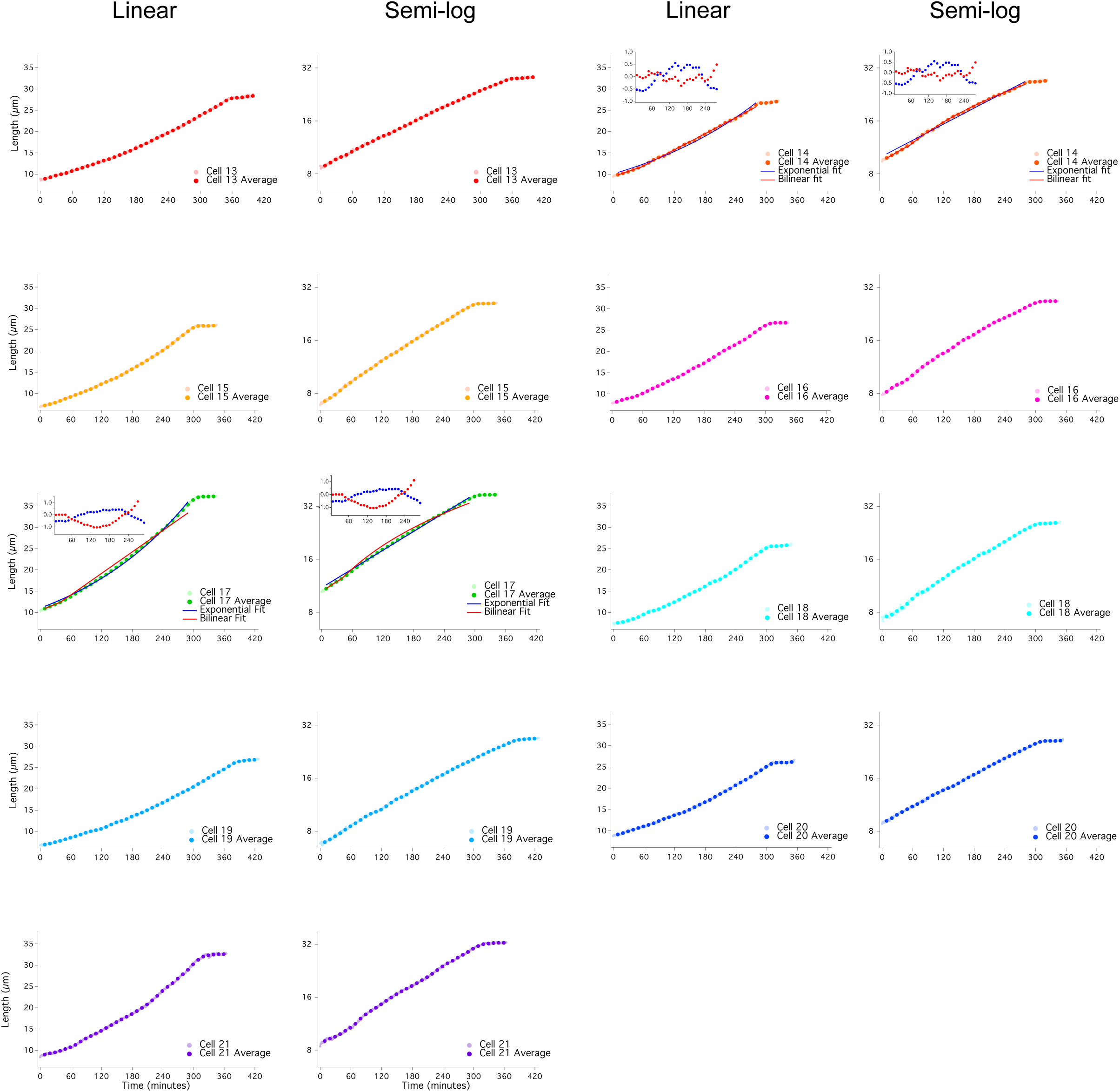
Individual growth data for 9 *ZEV:wee1* expressing cells. Lengths of nine individual *ZEV:wee1* expressing (yFS970) cells were recorded at 1 minute intervals from birth to septation following treatment with 31 nM β-estradiol and plotted on a linear or log scale. Cell length was averaged every 10 minutes in overlapping 20 minute windows. Fits from Figure 3 are included.

**Figure S4:**
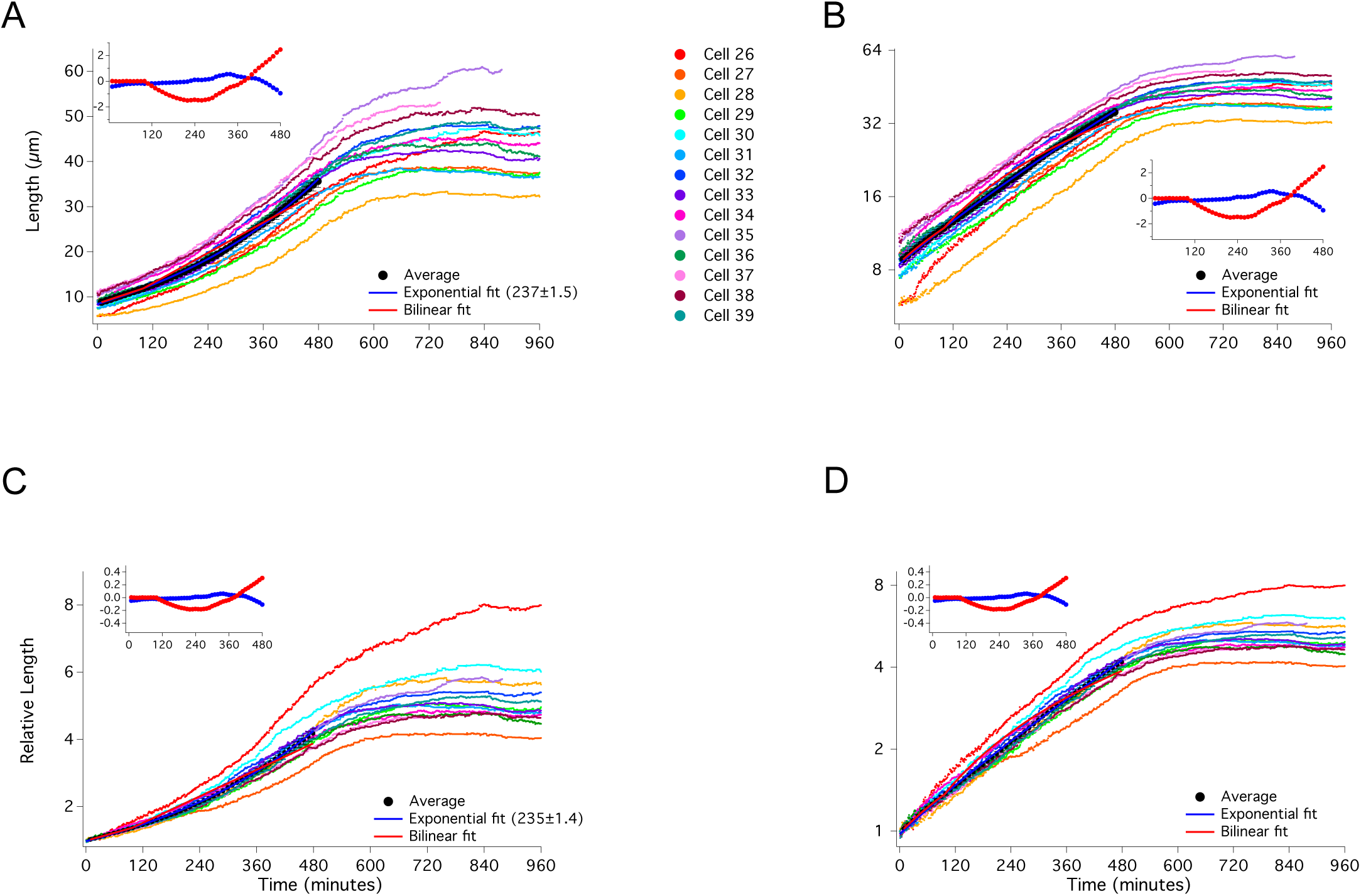
Full growth trajectories for *adh1:wee1* cells. Lengths of fourteen *adh1:wee1-50* (yFS145) cells were recorded at 1 minute intervals for 16 hours following a shift from 35°C to 25°C and plotted on a linear (A) or log (B) scale. The average of the cells in 10 minute windows ± standard error of the mean is shown. Exponential fit (with doubling time in parentheses), bilinear fit, and residual plot for these fits (inset) are shown. Birth lengths of cells were normalized to 1 and plotted using a linear scale (C) or a log-scale (D).

**Figure S5:**
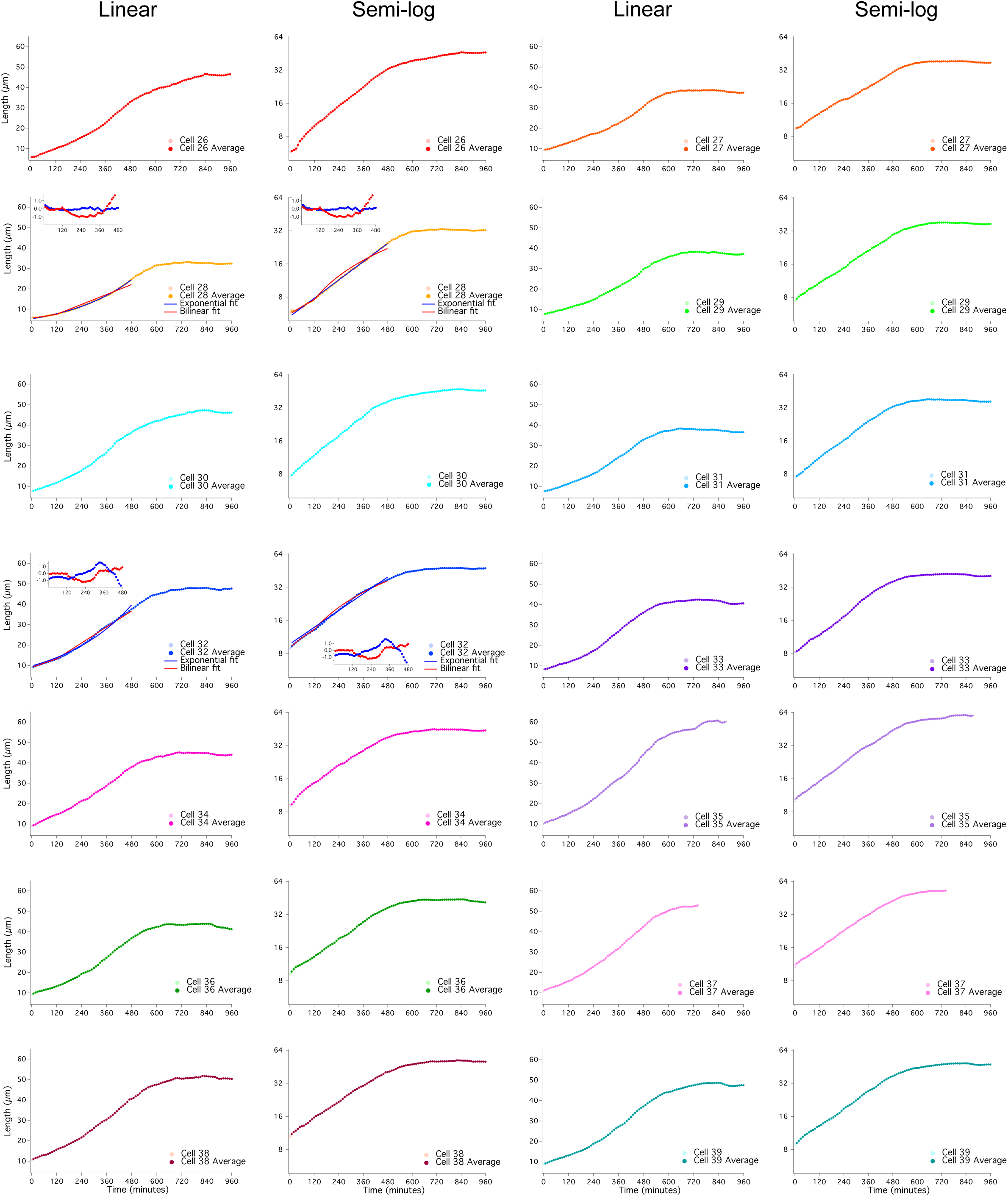
Individual growth data for 14 *adh1:wee1-50* expressing cells. Lengths of fourteen *adh1:wee1-50* (yFS145) cells were recorded at 1 minute intervals for 16 hours following a shift from 35°C to 25°C and plotted on a linear or log scale. Cell length was averaged every 10 minutes in overlapping 20 minute windows. Fits from Figure 4 are included.

